# Excitable dynamics of flares and relapses in autoimmune diseases

**DOI:** 10.1101/2023.05.03.539265

**Authors:** Yael Lebel, Tomer Milo, Alon Bar, Avi Mayo, Uri Alon

## Abstract

Many autoimmune diseases show flares in which symptoms erupt and then decline. A prominent example is multiple sclerosis (MS) in its relapsing-remitting phase. Mathematical models attempting to capture the flares in multiple sclerosis have often been oscillatory in nature, assuming a regular pattern of symptom flare-ups and remissions. However, this fails to account for the non-periodic nature of flares, which can appear at seemingly random intervals. Here we propose that flares resemble excitable dynamics triggered by stochastic events and show that a minimal mathematical model of autoimmune cells and inhibitory regulatory cells can provide such excitability. In our model, autoimmune response releases antigens that cause autoimmune cells to expand in a positive feedback loop, while regulatory cells inhibit the autoimmune cells in a negative feedback loop. The model can quantitatively explain the decline of MS relapses during pregnancy and their postpartum surge based on lymphocyte dynamics, as well as the decline in MS relapses with age. The model also points to potential therapeutic targets and predicts that even small modulation of regulatory T cell production, removal or activity can have a large effect on relapse rate. Excitable dynamics may underlie flares and relapses found across autoimmune diseases, thus providing an understanding that may help improve treatment strategies.

## Introduction

Autoimmune diseases are disorders in which the immune system attacks healthy cells and tissues. These conditions can affect many different organ systems with diverse symptoms. One common feature of many autoimmune diseases is the occurrence of flares, also called relapses ^1^. Relapses are episodes of increased disease severity. Many autoimmune diseases are thus characterized by relapses and remissions in which symptoms appear and then subside. Examples include multiple sclerosis (MS) ^2^, rheumatoid arthritis ^3^, inflammatory bowel disease^4^ (IBD), lupus ^5^, myasthenia gravis^6^ and psoriasis^7^.

Several factors can trigger a flare, including infections, stress and exposure to certain drugs and other environmental factors ^8–10^. It is believed that these triggering factors may disrupt the balance between immune effector cells and regulatory cells, causing an enhanced autoimmune attack ^11,12^.

The rate of relapses varies between autoimmune diseases. In MS, one of the best studied diseases, relapse rate averages about 0.6/year ^13^.

The development of autoimmune diseases and flares is influenced by genetic and environmental factors, but the specific mechanisms underlying these processes are not yet fully understood^14^. A better understanding of the underlying mechanisms that contribute to flares could inform treatment strategies aimed at reducing relapse rates. Therefore, it is important to identify potential unifying mechanisms that could explain the occurrence of flares across different autoimmune diseases.

Studies on multiple sclerosis have focused on models of the immune system that give rise to oscillatory flare dynamics ^15–17^. Flares in these models thus have a defined period. However, this periodicity is at odds with clinical data in which relapses do not seem to have a defined period ^13,18^. Moreover, these models typically do not incorporate the impact of environmental triggers on flare occurrence, which have been suggested to be major contributing factors to flare events in autoimmune diseases ^19,20^.

Here we hypothesize that flares result from excitable dynamics in the immune system. Excitability is a property in which a triggering event causes a large pulse of activity that returns to a locally stable baseline. Unlike oscillatory models, each new pulse requires a new triggering event, and flares thus appear stochastically with no defined period. We present a minimal model of immune effector and regulatory cells which shows excitability. Flares are triggered by factors such as stress and infection that enhance immune activity beyond a threshold. We compare the model predictions to large datasets on MS relapses and describe relapse changes during pregnancy. The model suggests that mild interventions in key parameters can profoundly reduce the relapse rate, offering directions for future treatment strategies.

## Results

### Relapse statistics in MS suggest stochastic excitability rather than oscillatory dynamics

To investigate the frequency of relapses we analyzed the Multiple Sclerosis Outcome Assessments Consortium (MSOAC) database^21^. This database contains records of patients in placebo arms of clinical trials done in MS patients from the year 2012 onwards. The database contains 139 relapses with exact dates from 31 patients with relapsing-remitting MS. A time course for one patient is shown in Fig 1A. The relapse-free survival probability (see methods) is exponential, with a mean of 191±10 days (Fig 1B). In other words, the probability of a relapse per unit time, known as the relapse hazard, is roughly constant in time.

**Fig 1:**
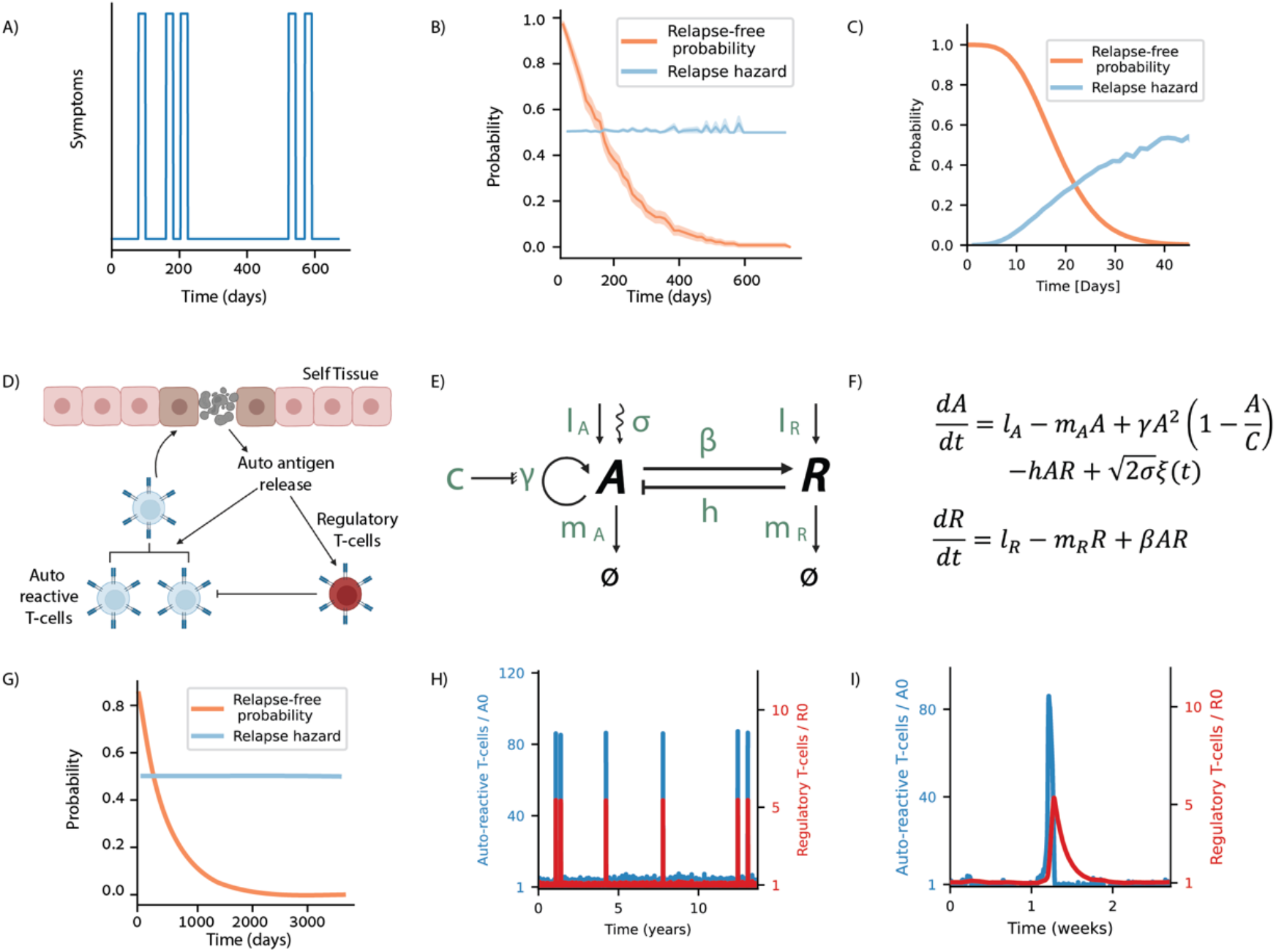
Excitable model of the immune system shows stochastic flares. A) A time course of a patients’ symptoms taken from the MSOAC database. B) Relapse-free times and relapse hazard rates of all patients in the database with precise relapse dates. C) Relapse-free survival and hazard for oscillatory behavior from 500 simulations. D) Schematic of the relapse model in which autoimmune effector cells A kill tissue cells, releasing autoantigen that enhances proliferation of A and of regulatory cells R that inhibit A. E) Circuit diagram of relapse mode. F) Model Equations. G) Relapse-free survival and hazard in the relapse model from 1000 simulations. H) Stochastic simulations show spike-like flares of A. I) Focus on a single flare with a rise in A followed by a rise in R, a decline in A and a slower decline of R. Fig 1D was Created with BioRender.com.

This analysis precludes an oscillatory mechanism of relapses, which would have a specific inter-relapse time, so that the relapse free survival drops around a specific interval duration (Fig 1C). The observed exponential distribution of relapse-free survival times does not have such a typical interval time. This observation, together with the spike-like nature of the relapses, suggest a mechanism in which stochastic effects cause flares in an excitable system.

### An excitable model of the immune system shows stochastic flares

We developed a minimal mathematical model of autoimmune flares based on well characterized immune interactions. Our goal was to investigate whether a simple model of immune activity could exhibit excitability, characterized by the ability to produce stochastically occurring flares of activation with a constant probability per unit time, and a refractory period.

We focus on autoimmune effector cells, which we denote as *A*. These CD4 and CD8 T cells can target and attack healthy tissue (Fig 1D). This attack releases self-antigens, which raise the proliferation rate of the effector cells ^14,22^. The autoantigens also induce proliferation of regulatory cells, such as Foxp3+ Tregs, which we denote as *R* ^23^. These regulatory cells suppress the effector cells ^14,24^.

We model these interactions using differential equations for the rate of change of A and R (Fig 1D, E). We explored various mathematical models based on the known interactions between these two cell types and ultimately arrived at a simple model that exhibited strong excitability (see SI for alternative models).

This model, which we call the *relapse model*, is based on the production and removal of effector cells (A), whose proliferation rate is proportional to autoantigen levels. As autoimmune response releases antigen, the autoantigen level is also proportional to A, resulting in an autocatalytic proliferation of A that goes as *A*^2^. To avoid singularities in which A goes to infinity, we employ a carrying capacity term in which proliferation drops to zero when A reaches a maximal level C ^25,26^. This carrying capacity can be due to the limited number of naive cells with the required reactivity, their maximal clone size determined in the lymph node ^27^, and limitations of physical space and growth factors.

The effector cells are inhibited by regulatory cells R as described by a negative term proportional to the product AR. This term models effects which require proximity of A and R cells, including direct killing by contact or local consumption of IL2 by R cells ^28,29^. Under normal conditions, regulatory cells are produced and eliminated at baseline levels, meaning that the number of cells in each lymph node remains relatively stable. However, in the presence of autoantigen, their replication rate increases, leading to an increase in the number of regulatory cells, represented by a term proportional to AR. Our model assumes that the maintenance of tolerance towards autoreactive T cells at baseline levels is mainly driven by the baseline levels of regulatory T cells, rather than the natural turnover of these cells ^30^.

To model stochastic activation of the effector cells we add a noise term to the equation for A. The noise term is essential to trigger the flares. The noise term can represent at least two known factors that enhance the rate of relapses. The first is infections which can activate autoimmune responses in prone individuals ^31^. This is effectively an increase in A production rate. The second is stress, which acts through endocrine and neuronal pathways to affect the immune system ^32,33^. We note that a noise term can also be added to the R equation, although this does not result in qualitative changes to our conclusions, and so we retain a noise term only in the A equations [SI].

Simulations of the resulting differential equations show stochastic spikes (Fig 1H). In each spike, A rises sharply, followed by a rise in R which inhibits A (Fig 1I). The spike ends with an exponential decrease of R. A new spike is formed when noise triggers the excitable system again. The relapse model shows flares whose timing is exponentially distributed (Fig 1G), as observed.

The dynamics produced by the relapse model qualitatively agree with studies that show an increase of T helper cells during relapses and an enhanced presence of regulatory cells during remission ^34,35^.

Our findings suggest that the relapse model, which captures key immunological interactions, can exhibit excitability and generate flares or relapses that resemble those observed in MS and other autoimmune diseases.

### Rate and shape of flares are governed by specific model parameters

To understand the relapse model, we employ a phase portrait analysis using nullclines ^36^ (Fig 2A). Each nullcline represents the steady state of one variable when holding the other variable fixed. This decomposes the feedback loop into two arms ^37,38^ : one arm in which A induces R, and the other arm in which R inhibits A. The crossing points of the two nullcline curves are the fixed points of the system.

**Fig 2:**
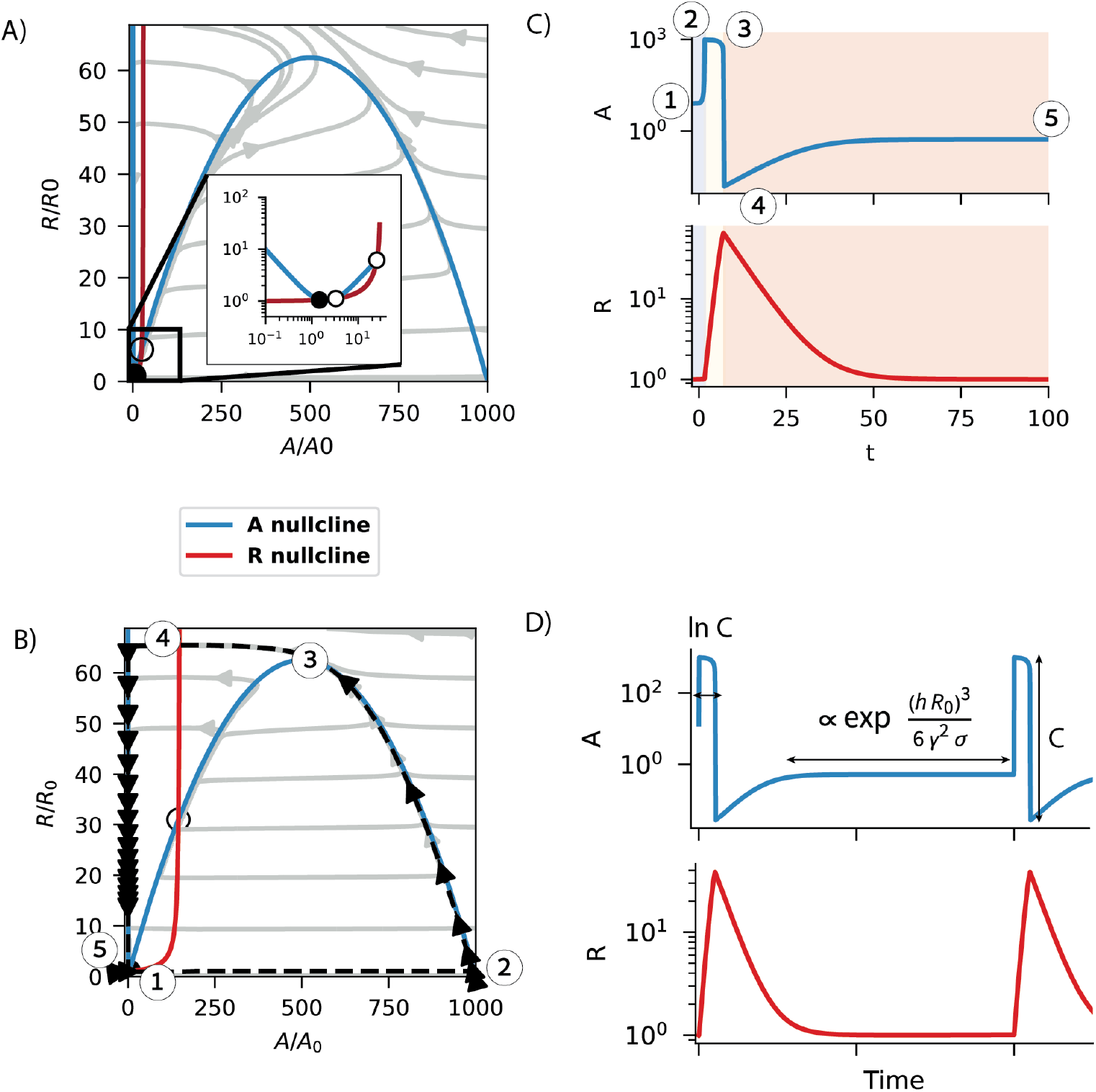
Analysis of the noise-induced flare. A) Phase portrait of the excitable system. B) Relapse trajectory in phase space. C) Time evolution of both autoimmune and regulatory T-cells during a flare. D) Parameter dependency of the different parts of the time evolution. Model parameters are: *B* = 0.001, *G* = 0.25, *D* = 0.15, *C* = 1000, in units where flare duration is about two weeks, and flare rate is about 1/year [Methods].

The first nullcline (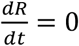, in red) shows how the R steady state monotonically rises with A. The second nullcline (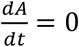, in blue) has an inverse-N shape. Inhibition of A by R explains the leftmost descending arm of the nullcline. When R is at moderate levels, auto stimulation of A dominates over its inhibition by R, which results in an ascending curve. Finally, at high A, effector cell production decreases as the population approaches its carrying capacity, resulting in another descending arm of the nullcline. The inverse-N shape of the A nullcline is important for excitability. Excitability occurs when the nullclines cross in the first declining segment of the N-shaped nullcline. This provides a single stable fixed point, representing a physiological state in which autoimmune effector cells are present and held in check by regulatory cells. The presence of such autoimmune T cells in the healthy population which can target organs involved in autoimmune diseases such as MS, type-1 diabetes and thyroiditis, has been documented ^30^.

When noise-namely infection or stress - pushes the system away from the stable fixed point and across the ascending branch of the *A* nullcline, the system enters the excitable region (point 1 in Fig 2B). Effector cells *A* grow rapidly due to the A^2^ term and reach a value close to their carrying capacity C (point 2). This induces growth of R cells (point 3) which inhibits the *A* cells until they return to the first descending branch of the *A* nullcline (point 4). Finally, *R* cells decline, and the system returns to baseline (point 5).

The dynamics of a flare is shown in Fig 2B, C, with the key timepoints marked for clarity.

Noise triggers flares by pushing the system across a threshold. To calculate the rate of flares analytically we used the Kramer approach. This method involves calculating an effective potential that determines how likely the system is to cross a threshold after being perturbed by noise. The potential difference, or *ΔU*, represents the potential barrier that the system must overcome in order to start a flare - a large excursion in the phase space that returns to the fixed point. The probability of the system crossing this potential barrier is proportional to 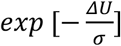, where σ is the amplitude of the noise. In other words, the larger the potential difference and the smaller the noise, the less likely the system is to cross the barrier and generate a relapse. The effective potential and potential difference are therefore key factors that determine the rate and shape of flares in our model. An exact solution is provided in Methods.

This barrier-crossing dynamics produces an exponentially distributed flare spacings, with mean flare rate:

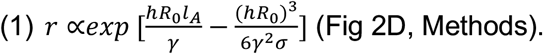

The standard deviation of the flare timing is equal to the mean since the flare timing distribution is exponential.

The model parameters also affect the shape of each flare (see methods - Relapse Shape and Duration for details). The flare amplitude is proportional to the effector cell carrying capacity C, whereas flare duration is proportionate to *ln C*, due to the super-exponential growth of A during flare (Fig 2C). After the flare, effector cells drop below baseline and regulatory cells are above baseline, making it unlikely to have another relapse. This is a refractory period, whose duration is proportional to 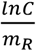. The refractory period is typically much shorter than the inter-spike time, and thus the exponential relapse-free survival distribution is a good approximation at times longer than the refractory period.

We provide a table of estimated parameters in terms of dimensionless parameter groups (Table 2). T cell lifetimes (both effector and regulatory) are on the order of days. Since flares entail sufficient damage to cause symptoms, we assume a carrying capacity a thousand times larger than baseline, C=10^3^ to allow for large spikes^39^ - the results remain qualitatively the same for values of C that are C=10 or larger. In the simulations we use dimensionless effector cell reactivity 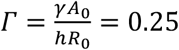 where the steady state is *A*_0_ and *R*_0_ for *A* and *R* cells respectively.

**Table 1:** model parameters.

| Parameter | Meaning | Units |
| --- | --- | --- |
| $l_A$ | Rate of production of auto-reactive cells | cell/time |
| $l_R$ | Rate of production of regulatory cells | cell/time |
| $m_A$ | Natural turnover time of auto reactive cells | 1/time |
| $m_R$ | Natural turnover time of regulatory cells | 1/time |
| $\gamma$ | Reactivity of auto reactive cells to antigen | (cell · time)<sup>-1</sup> |
| $C$ | Carrying capacity of auto reactive cells | cell |
| $h$ | Rate of inhibition of auto reactive cell by regulatory cells | (cell · time)<sup>-1</sup> |
| $\beta$ | Reactivity of regulatory cells to antigen | (cell · time)<sup>-1</sup> |
| $\sigma$ | Noise amplitude | cell<sup>2</sup>/time |

**Table 2:**
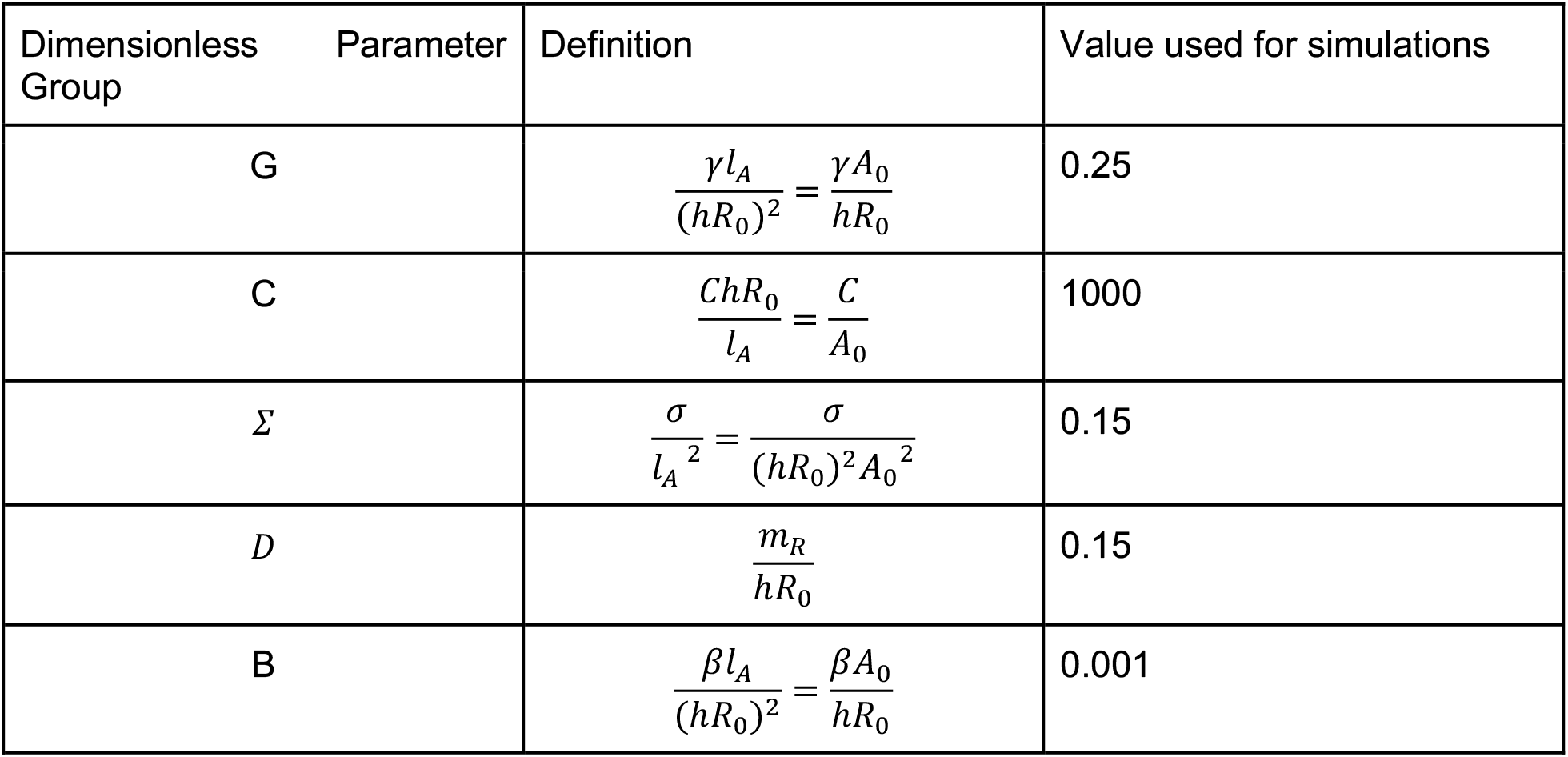
Dimensionless parameters values used for simulations.

The range of parameters that provides excitability is large (Fig 3A)-a range of several orders of magnitude foreach parameter around the parameters used for Fig 2A-C-as described in more detail in the next section.

**Fig 3:**
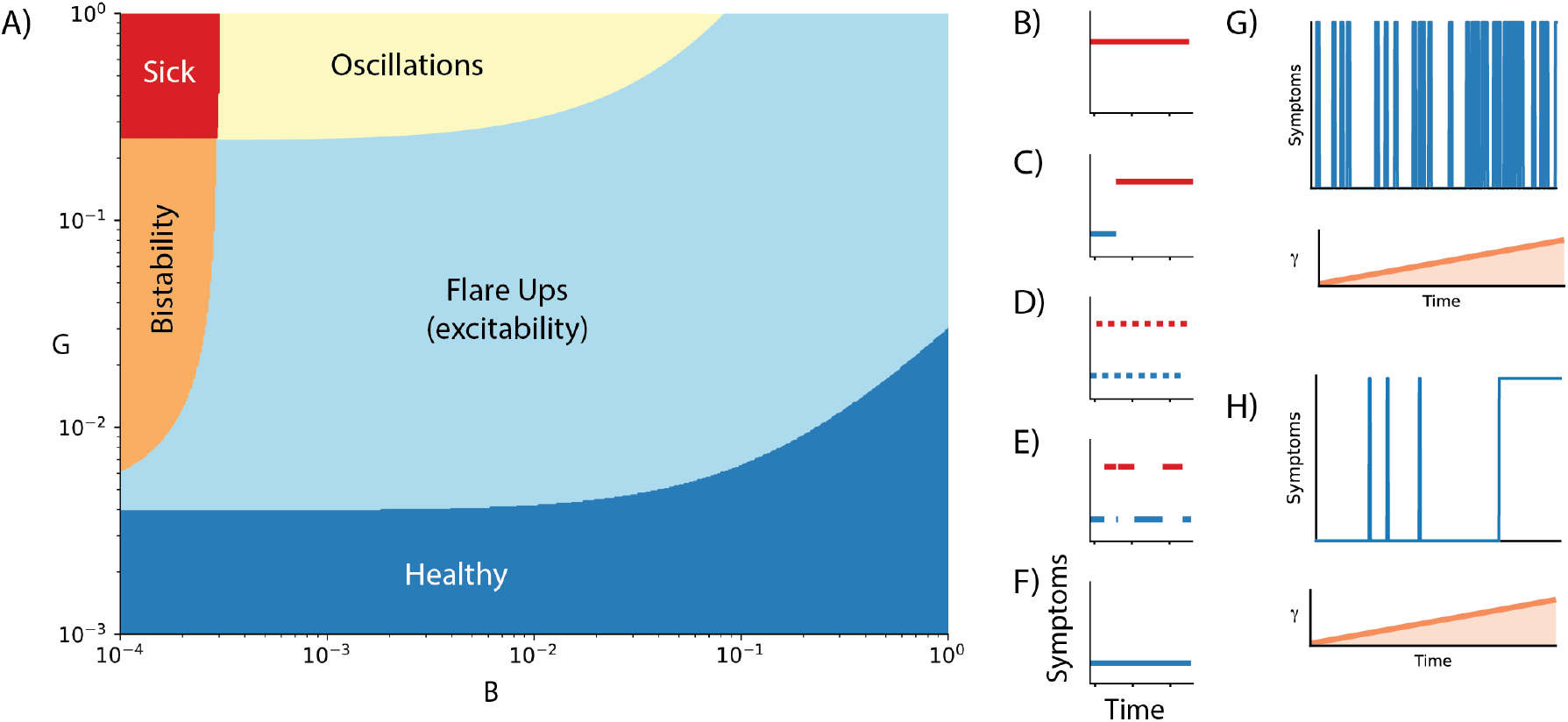
Phase diagram of the model and transitions to progressive disease. A) Phase diagram of the different regimes of behavior as a function of the auto-stimulatory parameter *γ* and regulatory stimulation parameter *β*. B-F) Qualitative time series for the different regimes: B) Single sick state. C) Bistability of the healthy and sick states. D) Oscillations between healthy and sick states. E) Flare ups. F) Single healthy state. G) simulations with increasing gamma showing onset of excitable flares which transition to rapid oscillations. H) simulations with increasing gamma showing onset of excitable flares which transition to chronic activation. Dimensionless parameters (see Methods): *G*(*τ*) = 0.25 + 5 ⋅ 10^−6^*τ, B* = 5 ⋅ 10^−3^ in G, *G*(*τ*) = 0.03 + 1.4 ⋅ 10^−5^*τ, B* = 3 ⋅ 10^−4^ in H.

Notably, the model exhibits strong flares even though autoimmune cells and regulatory cells have similar turnover rates. This is unlike standard models of excitable systems such as the Fitzhugh-Nagumo model which describes, for example, neuronal spikes. These standard models depend on separation of turnover times where the fast variable can rise before the slow inhibitory variable can overtake it. The reason that the present model shows strong flares is the auto stimulation in the *A* equation 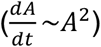. This causes a super-exponential rise in autoimmune cells - a finite time singularity which is prevented by their carrying capacity. This super-exponential rise outruns the rise in regulatory cells. The stark nature of the flare can be seen in the nearly horizontal dynamical arrows in figure 2A, which effectively creates a separation of timescales.

### Transitions between flares and chronic disease

MS patients can transition from a relapse-remitting disease to a more chronic form called secondary progressive disease in which disability becomes progressively worse. To understand such a transition we consider the possible behaviors of the model when parameters change.

We depict these behaviors in a phase diagram in Fig 3A. Excitability occurs in a wide region of parameters. Regimes other than excitability occur when parameter thresholds are crossed. When the autoantigen stimulation parameter *γ* is very low the model shows a monostable situation with no flares - corresponding to a healthy state. When *γ* is very high the model shows a chronically sick state with constant high autoimmune activation - similar to chronic autoimmune disease states. A region of bistability occurs when regulatory T cells are weakly triggered by antigen (low *β*) - this effectively corresponds to a chronic autoimmune disease once triggered. Finally, oscillations (instead of excitability) occur in a region of parameter space with high *γ* and intermediate *β* (Fig 3).

One pathway from health to flares to chronic state may be a gradual increase in the stimulation parameter *γ*. This increase may be due to epitope spreading in which damage to a tissue causes the establishment of effector cells (T cells and B cells) to a wider range of antigens than before the damage ^40–42^. In principle, every flare has a chance to increase *γ*, making the system more auto stimulatory. A trajectory of increasing *γ* can transition between the healthy part of the phase diagram, through the excitable region, and finally to a bistable or oscillatory region (depending on *β*) (Fig 3G, H). In the oscillatory regime, the spikes of the oscillation occur at a period determined by the refractory period, and thus may resemble a chronic autoimmune attack in which each spike is followed closely by another. Both the bistable and oscillatory endpoints may thus point to a progressive increase in disability.

### Model predicts the changes in relapse rate in pregnancy and postpartum

To test the model, we consider the changes in autoimmune relapse rates in pregnancy and the postpartum period. During pregnancy, the immune system undergoes changes that suppress harmful responses to the fetus, leading to a decrease in the severity of some autoimmune disorders. This is due to an increase in regulatory T-cells and anti-inflammatory cytokines, as well as hormonal changes such as an increase in estrogen and progesterone.

Data on the relapse rate of MS shows a decline during pregnancy followed by a rebound in the first three months after delivery ^43^, Fig 4A.

**Figure 4:**
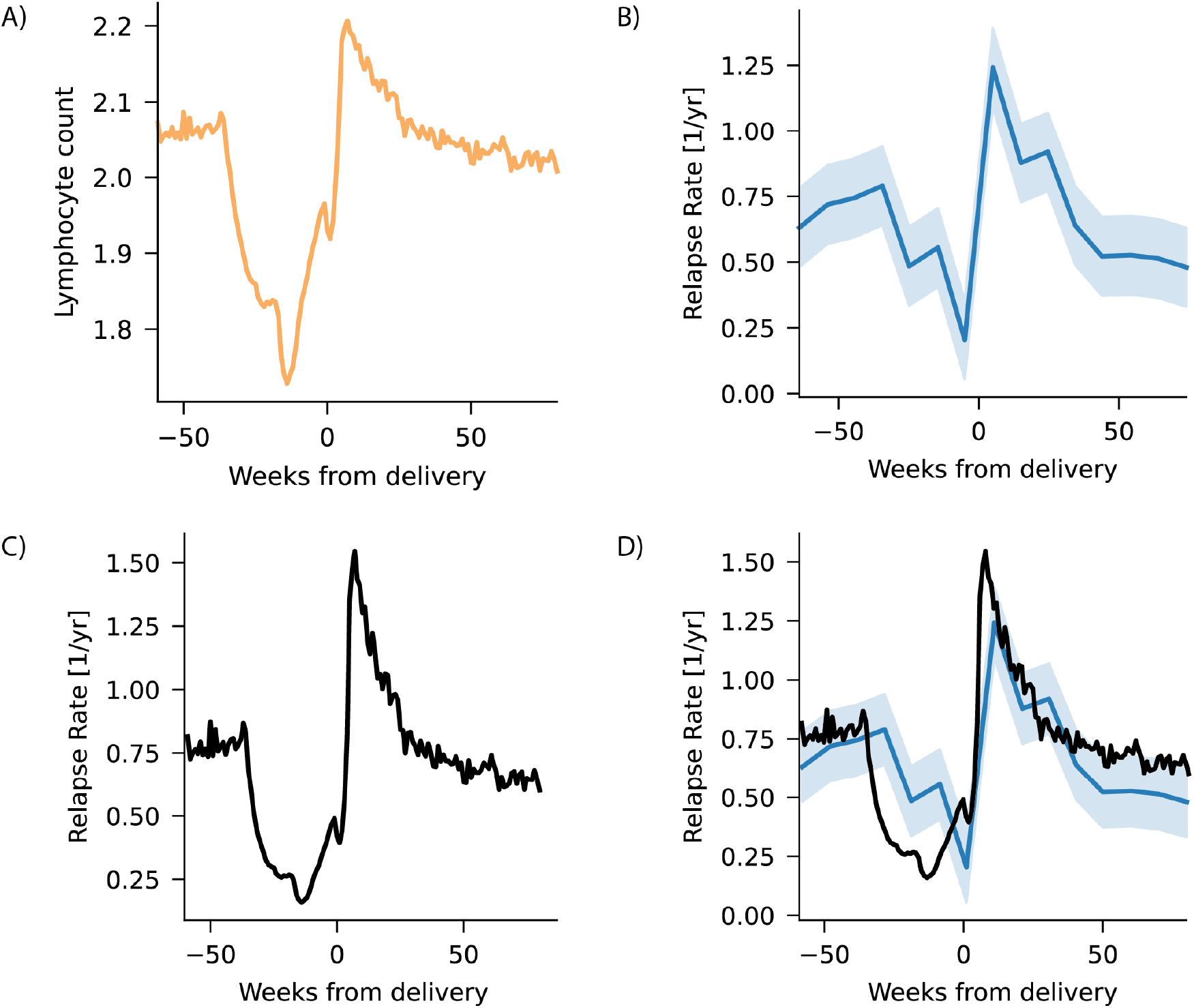
Model predicts MS relapse rate variation in pregnancy and postpartum based on lymphocyte counts. A) Lymphocyte count blood tests from the Clalit dataset (n=1.4 million tests) averaged over each week from 60 weeks before delivery to 80 weeks after. B) MS relapse rate averaged over three-month periods from Vukusic et al. C) Relapse rate computed from model Eq with *r*_*baseline*_ =0.6/year and *q* = 10. D) Overlay of computed and measured relapse rates.

**Fig 5:**
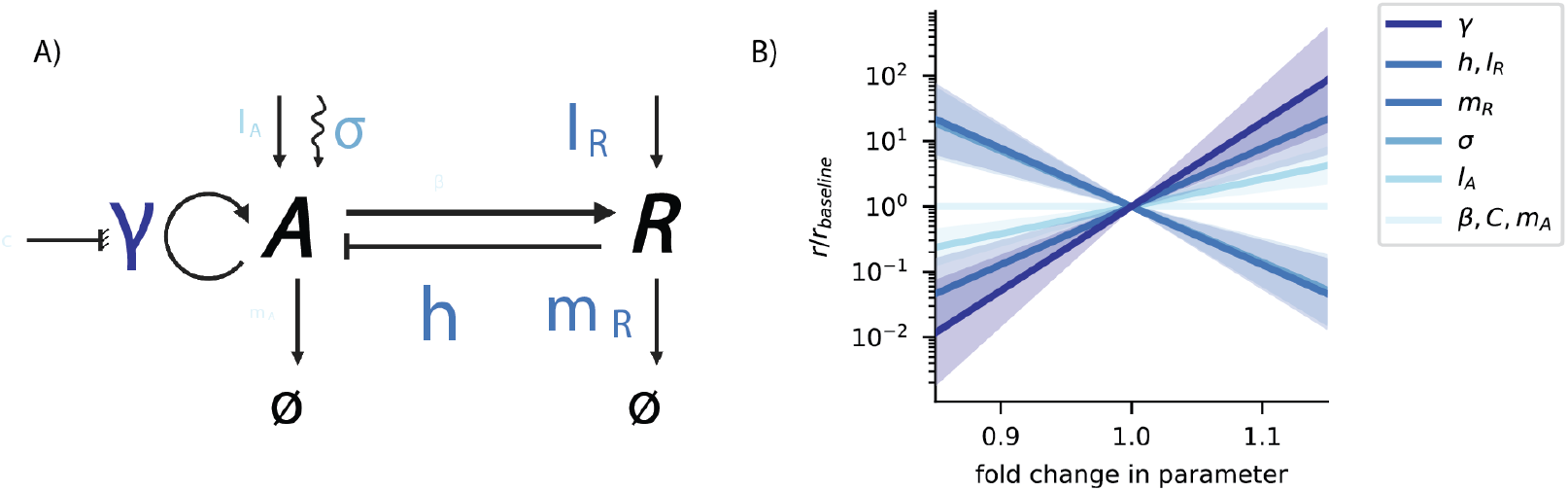
Certain parameter changes can strongly reduce the relapse rate. A) Parameters in the circuit, letter size indicates the sensitivity of relapse rate to each parameter. B) Relative change in relapse rate for a relative change in each parameter. Note the logarithmic y axis. Baseline parameter group is q=10.

To compare this to the model, we obtained data on the temporal dynamics of lymphocytes during pregnancy from the Clalit dataset on 300,000 pregnancies. This dataset includes 1.5 million lymphocyte counts averaged over each week of pregnancy and 80 weeks postpartum Fig 4B. Lymphocyte counts decrease during pregnancy and increase after delivery, returning to baseline (pre-pregnancy) level after 60 weeks.

We reasoned that lymphocyte counts can serve as an approximate measurement for *A*_0_, the baseline level of effector cells. We furthermore reasoned that the combined effect of decrease in total amount of lymphocyte count and the relative expansion of regulatory T-cells ^39^ can be approximately described by constant levels of *R*_0_. We then used equation (1) to calculate the relapse rate relative to baseline. The relation between relapse rate *r* and the relative change in lymphocyte levels *𝜖* can be written as *r* = *r*_*baseline*_*e*^*q𝜖*^ with a single fit parameter *q*, using *r*_*baseline*_ =0.6/year. We find that this equation captures the observed dynamics of the normalized relapse rate well with *q* = 10 ± 4 (correlation coefficient 0.82, p<0.005).

### Model suggests parameter modulations to reduce the relapse rate

To explore potential targets for intervention, we asked which parameters of the model affect the relapse rate most strongly. For this purpose, we evaluated the sensitivity with respect to each parameter in the mathematical model. Sensitivity is defined as the logarithmic derivative of the relapse rate with respect to the parameters [Methods], which is roughly equal to the percent change in relapse rate for a 1% change in the parameter. We used the estimated parameters from the pregnancy dataset as the baseline system parameters.

We found that most parameters had high sensitivity, with a 1% change in parameters resulting in a greater than 10% change in the relapse rate. The parameters with the largest effect were the baseline level of regulatory T-cells (*R*_0_), their inhibitory effect (*h*), and the autostimulation parameter *γ*.

Although the carrying capacity (C) and the activation rate of regulatory cells (*β*) are included in our model, they have negligible effect on the relapse rate as their values are significantly far from those of the other parameters. A third parameter had negligible effects in the assumed parameter range - the natural turnover rate of autoreactive cells *m*_*A*_, as it was assumed to be the less dominant timescale in the system. These parameters only affected the shape and amplitude of each relapse.

The high sensitivity for many parameters is optimistic for treatment strategies. Suppose that our aim is to reduce relapse rate by a factor of 100, which practically guarantees lack of relapses because it converts the ∼1/year relapse rate to less than 1/lifespan. To achieve this one need only have a 15% decrease of *γ*, or a 22% increase in either *l*_*R*_ or *h*. A combination of parameter changes would require even smaller effects, such as changing both *l*_*R*_ and *h* by 11%. Thus, our findings provide hope for developing effective treatments that target the immune system parameters related to the relapse rate of multiple sclerosis.

The immune system is a complex and delicate system that requires a fine-tuning of its parameters for proper function. While the model allows for a large variety of inter-relapse times, in reality, only a limited range of parameter combinations can produce relapse rates that are characteristic of an active disease, while still allowing for patient survivability. Therefore, the specific combination of parameters that is required for disease activity is likely to lie within a small region of the parameter space that is consistent with the fine-tuned parameters of the immune system.

## Discussion

We present an excitable mechanism for flares in autoimmune diseases based on a minimal model of the adaptive immune system. The model describes interactions between autoimmune cells and inhibitory regulatory cells and shows robust excitatory spikes that occur at random times. We present evidence that flares in MS are not oscillatory but rather have an exponential inter-flare time distribution, which is explained by the presented model. The model can explain the decline of MS relapse during pregnancy and the postpartum surge in relapses based on longitudinal lymphocyte measurements. It also predicts which interactions might serve as targets to reduce relapse rates. It predicts that mild intervention in regulatory T cell parameters can have large beneficial effects on reducing relapse rate, offering insight to potential treatment strategies.

The present model indicates that a relapse is triggered when autoimmune cell activation is raised beyond a threshold. The dynamics show a rapid rise in autoimmune activation, reaching close to its carrying capacity, followed by a rise in regulatory cell activation which shuts off the flare. Several factors may be responsible for triggering the flare, and these are described as noise in the model that affects autoimmune activation stochastically. One key factor is infections, which are known to often occur immediately before flares in autoimmune diseases such as MS ^44^, lupus ^5^, and other diseases. Other factors include stress, which has pleiotropic effects on the adaptive immune system, and which can incite autoimmune flares ^45,46^.

Excitable systems have been extensively studied in neurons, and their dynamics have been understood using canonical models like the Fitzhugh-Nagumo model ^47–49^. Typically, strong flares in these models depend on having a fast variable and a slow variable with strong separation of timescales ^50^. The present model provides strong flares even without explicit separation of timescales between the two equations, due to a strong autocatalysis of the autoimmune cells. This results in similar timescales near the stable fixed point, but very fast dynamics for A when the spike is triggered. Indeed, autoimmune cells and regulatory cells are expected to have similar turnover times.

The present model differs from previous mathematical models that treat autoimmune flares as oscillations, because it treats flares as an excitable system. Oscillator models predict a periodic appearance of flares, or if they include noise, a periodic appearance with some stochastic variation, whereas in reality flares usually do not seem to have a characteristic period, as we find in MS data. The present model predicts an exponential distribution of inter-flare times, since flares are triggered by a stochastic process with essentially constant probability per unit time.

We note that oscillatory dynamics in other (non-autoimmune) diseases have been documented^49^. The present model has a small region of parameter space in which it has oscillatory solutions.

The model is agnostic to the precise antigen and target tissue, and thus may potentially apply to a wide range of autoimmune diseases with flares; these include SLE (lupus), rheumatoid arthritis, inflammatory bowel syndrome and psoriasis.

One may speculate that additional diseases may have a currently unknown flare dynamics. One such disease is type 1 diabetes^51^. This disease has a ‘honeymoon phase’ in which initial treatment often causes glucose control to return for a few months, followed by relapse of the disease. This may indicate flare dynamics. It would be interesting to explore whether chronic diseases such as Hashimoto’s thyroiditis also have flare dynamics during their subclinical phase.

The model might provide insight into the transition from relapsing-remitting disease to chronic (secondary progressive) disease as occurs in MS. This transition may be due to slow changes in the model parameters that cross between the excitable regime and the bistable or chronic regime in which autoimmunity is constantly active. These slow changes in parameters may occur with the number and severity of relapses. For example, a rise in γ, the amplification of autoimmune cells, can cause a shift to chronic autoimmunity. This may occur due to increasing antigen-sensitivity that occurs with an increasing number of flares due, for example, to antigen spreading ^40–42^.

The model can potentially also address changes in autoimmune flares with age. Aging generally increases memory Tregs and decreases memory effector cells ^52^, together with declining total lymphocytes. The model therefore predicts a decline in relapse rate which is indeed observed in MS ^53^.

The model may be useful to better understand the effect of immunomodulatory treatments on autoimmune diseases. Immune checkpoint treatments for cancer, for example, impair Treg function and increase the risk of multiple autoimmune diseases^54^. Treatments for autoimmune diseases such as CD3 neutralizing antibodies ^55–57^ reduce effector cells and enhance regulatory T cells, and are predicted by the model to significantly reduce the relapse rate.

In summary, we presented a mechanism in which the interplay of autoimmune and regulatory cells can cause robust flares triggered stochastically by factors such as infection and stress. The model helps to quantitatively predict how changes in the adaptive immune system translate to changes in relapse frequency. It points to several key interactions as potential targets for reducing the relapse rate - even small modulation of regulatory T cell production, removal or activity is predicted to have strong effects on relapse rate. Such quantitative approaches can inform treatment strategies.

## Methods

### Relapse-free probability and relapse hazard curves from MSOAC database

The MSOAC database contains data on 4370 relapse events from 1151 different patients. The relapse events in the database are either entered with the exact start date of the event, or with the date it was recorded into the study. The latter can be later than the actual date of the event and depends strongly on the clinical visit frequency. To avoid this, we only used relapse entries that had an exact start date recorded. This amounts to 31 patients with 139 relapses total.

For each patient with exact relapse dates, we calculated the time differences between consequent relapses after the first relapse. We then calculated the cumulative probability of relapse-free periods. The relapse-free survivability is defined by *S*(*t*) = 1 − *CDF*(*t*), which is the probability to have a relapse in time shorter than t. The relapse hazard is defined as the probability per unit time of a relapse given by 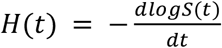.

### Model equations

The dynamics described above was modeled by two variables - the auto-reactive T-cells (*A*) and the regulatory T-cells (*R*). The dynamics can be described by:

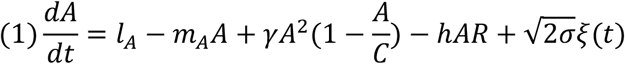

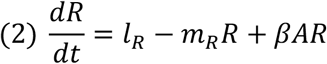

The parameters are described in the following table:

We assumed that the main mechanism of immune tolerance is regulatory T cells rather than the natural turnover of the autoreactive effector T cells, which means *m*_*A*_ ≪ *hR*_0_. This sets a typical timescale for the system 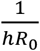.

Using dimensional analysis, we reduced the model to a dimensionless model with five parameters:

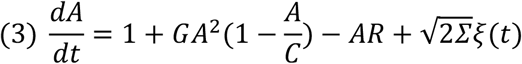

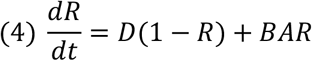

where now the rescaled auto-reactive T-cells are in units of their baseline levels 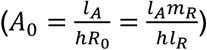, the regulatory T-cells are in units of their baseline levels 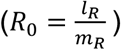 and time is in units of 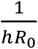. We assumed the turnover time of the regulatory cells to be identical to that of the auto-reactive cells. Other assumptions used were *B* ≪ 1 (this is necessary in order to have effective shut down of the flare) as well as dimensionless carrying capacity C much larger than all the other dimensionless parameters.

We modeled noise in the system (for example, viral infections and stress) by Gaussian white noise with mean zero and standard deviation *σ*. Adding a white nose term to Eq 4 does not change the conclusions, and thus for simplicity we added noise only to the effector cells in Eq 3 [SI]. The dimensionless parameters used for the simulations in the main text are shown in table 2.

### First passage time

In order to calculate the average time between flares, we used the Kramer approach ^58^, under the assumption that as long the system is not in a flare, the regulatory T-cells are at steady state (*R=constant*). This allows us to calculate an effective potential for the effector cell equation:

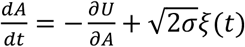

Since we assume that the steady state of both *A* and *R* is much smaller than C, namely *A*/*C* ≪ 1, the effective potential is:

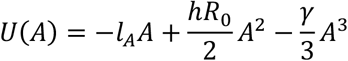

The potential *U*(*A*) has a minimum at the steady state *A*_*st*_ and a maximum at *A*_*th*_, above which the system enters the excitable regime. These points are the zeros of the derivative 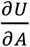:

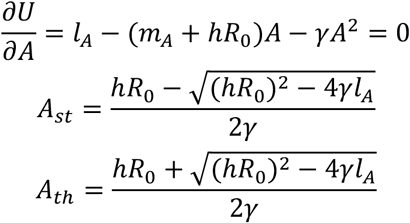

In order to calculate the mean crossing time, we need to calculate the potential difference between these two values. This results in 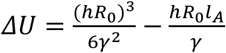, where we used the approximation *γl*_*A*_ ≪ (*hR*_0_)^2^. Using Kramer’s approximation ^58^ the average time to cross the threshold is:

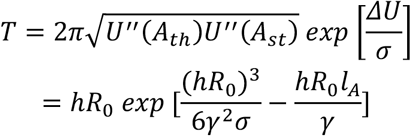

### Relapse Shape and Duration

Once a flare is initiated, the dynamics is dominated mostly by the deterministic part of the model. The flare can be divided into three parts, as is seen in Fig 2C. We will now analyze the duration of each of the parts:

1. Fast exacerbation: this regime is characterized by a quick rise to the carrying capacity C. The dominant part in the *A* equation is the quadratic and we assume *A* ≪ *C* for most of the way (omitting this assumption has no effect on the result - not shown). R remains in its steady state to a good approximation. We integrate the equation for the time evolution of A to find:

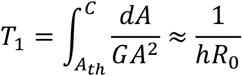
2. Shutdown of the flare by increase of the regulatory T cells. The trajectory in phase space slides along the A nullcline (which means 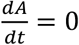, can be seen in Fig 2B) until 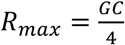. Along the nullcline, as *A*∼*C*, we can approximate:

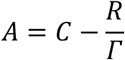
3. Inserting this into the time evolution for R and integrating we have:

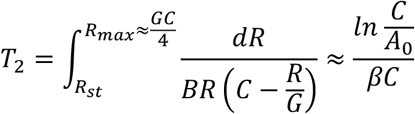
4. Slow shutdown of the regulatory response -refractory period. This regime is characterized by a slow decline along the declining leg of the A nullcline until returning to the steady state (see figure 2b). At this part of the nullcline we can approximate:

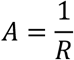

Inserting this into the time evolution of R and integrating we get:

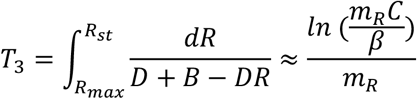

### Pregnancy

Using the expression for the first passage time, we obtain an expression for the relapse rate: 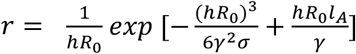. Using this, we estimate the effect that a small change of production rate *l*_*A*_→ (1 + *𝜖*)*l*_*A*_ has on the relapse rate: *r*(*𝜖*) = *r*_*baseline*_ *exp* [*q𝜖*], with 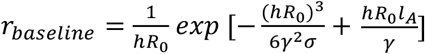 and 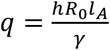. The baseline level was taken to be *r*_*baseline*_ = 0.64 which is the baseline (pre-pregnancy) relapse rate in Ref ^43^ in agreement with the average relapse rate known more widely from the literature. In order to compute α we used lymphocyte count of pregnant women from the Clalit database. To compute *α*, we divided the baseline level by the lymphocyte count of each week. The relapse rates used were extracted from^43^. Since the lymphocyte count is provided by weeks, we averaged over 13-week periods to match the relapse data measured in trimesters. The fit was done using Levenberg-Marquardt algorithm through Scipy’s curve_fit() tool, for a single fit parameter *q*.

### Sensitivity analysis

To perform sensitivity analysis, we assume a small change in the parameters *𝜖* ≪ 1, and take the linear approximations resulting, e.g for h:

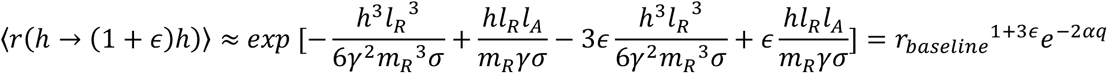

And a similar calculation for each of the other parameters. We used the value of *q* from the pregnancy data fit and a baseline level *r*_*baseline*_ = 0.64.

## Acknowledgements

This project was funded by the European Research Council (ERC) under the European Union’s Horizon 2020 research and innovation program (grant agreement No 856487). Data acquisition was approved by the Clalit Helsinki committee RMC-1059-20. The authors thank the investigators from the Multiple Sclerosis Outcome Assessment Consortium (MSOAC) Placebo Database for providing the data used in this study.

