## Supplementary information for "Excitable dynamics of flares and relapses in autoimmune diseases"

March 29, 2023

### 1 Detailed Mathematical and Dynamical Analysis of the Model Presented

#### 1.1 Model Equations

The model describes the dynamics of two variables: the auto immune effector or conventional T-cells (A) and the regulatory T-cells (R). The equations are:

$$\begin{aligned}\frac{dA}{dt} &= l_A - m_A A + a(A) A \left(1 - \frac{A}{C}\right) - hAR \\ \frac{dR}{dt} &= l_R - m_R R + a(A) R\end{aligned}$$

explanation of the terms

- Autoimmune cells are generated in the thymus at rate $l_A$ and are removed at rate $m_A$.
- Autoimmune cells are activated and proliferate as a response to a signal of antigen $a(A)$, up to a carrying capacity $C$.
- Autoimmune cells are removed by regulatory T-cells in a contact-based manner, i.e proportional to $-AR$, at a rate $h$.
- Regulatory T cells are generated in the thymus at rate $l_R$ and are removed at rate $m_R$.
- Regulatory T cells are activated and proliferate as a response to antigen signal $a(A)$.
- The antigen levels in this case are auto-antigens. Since autoimmune cells destroy tissue, they generate self-antigen . Thus the antigen levels are proportional to the amount of autoimmune cells $a(A) \propto A$.

Therefore the full equations are:

$$\begin{aligned}\frac{dA}{dt} &= l_A - m_A A + \gamma A^2 \left(1 - \frac{A}{C}\right) - hAR + \sqrt{2\sigma}\xi \\ \frac{dR}{dt} &= l_R - m_R R + \beta AR\end{aligned}$$

#### 1.2 Dimensionless Model

We now rescale the parameters to dimensionless form:

$$\tilde{A}=\frac{A}{A_0},\tilde{R}=\frac{R}{R_0}$$

$$A=A_0\tilde{A},R=R_0\tilde{R}$$

$$\frac{dA}{dt}=\frac{dA}{d\tilde{A}}\frac{d\tilde{A}}{dt}=A_0\frac{d\tilde{A}}{dt},\frac{dR}{dt}=R_0\frac{d\tilde{R}}{dt}$$

($A_0, R_0$ will be defined soon).

$$\begin{aligned}\frac{d\tilde{A}}{dt} &= \frac{l_A}{A_0}-m_A\tilde{A}+\gamma A_0\tilde{A}^2\left(1-\frac{A_0\tilde{A}}{C}\right)-h\tilde{A}R_0\tilde{R}+\frac{\sqrt{2\sigma}}{A_0}\xi \\ \frac{d\tilde{R}}{dt} &= \frac{l_R}{R_0}-m_R\tilde{R}+\beta A_0\tilde{A}\tilde{R}\end{aligned}$$

rescaling time:

$$\tau=Xt$$

$$\frac{d\tilde{A}}{dt}=\frac{d\tilde{A}}{d\tau}\frac{d\tau}{dt}=X\frac{d\tilde{A}}{d\tau},\frac{d\tilde{R}}{dt}=X\frac{d\tilde{R}}{d\tau}$$

(X will be defined). This results in:

$$\begin{aligned}\frac{d\tilde{A}}{d\tau} &= \frac{l_A}{XA_0}-\frac{m_A}{X}\tilde{A}+\frac{\gamma A_0}{X}\tilde{A}^2\left(1-\frac{A_0\tilde{A}}{C}\right)-\frac{hR_0}{X}\tilde{A}\tilde{R}+\frac{\sqrt{2\sigma}}{A_0X}\xi \\ \frac{d\tilde{R}}{d\tau} &= \frac{l_R}{XR_0}-\frac{m_R}{X}\tilde{R}+\frac{\beta A_0}{X}\tilde{A}\tilde{R}\end{aligned}$$

The steady state of A (i.e, with negligible autostimluation, or alternatively at $A\rightarrow 0$) is:

$$\begin{aligned}\frac{dA}{dt} &= l_A-(m_A+hR_0)\,A=0 \\ A_0 &= \frac{l_A}{m_A+hR_0}\end{aligned}$$

We assume that the main mechanism that preserves tolrance is the presence of regulatory cells rather than the natural turnover of effector cells, i.e $m_A \ll hR_0$, so the steady state ($A_0$) will be:

$$A_0=\frac{l_A}{hR_0}$$

this impies that the natural timescale of the equation is $hR_0$, which means we should take $X=hR_0$. Now inserting this in **??**:

$$\begin{aligned}\frac{d\tilde{A}}{d\tau} &= \frac{l_A}{hR_0\frac{l_A}{hR_0}}-\frac{m_A}{hR_0}\tilde{A}+\frac{\gamma A_0}{hR_0}\tilde{A}^2\left(1-\frac{A_0\tilde{A}}{C}\right)-\frac{hR_0}{hR_0}\tilde{A}\tilde{R}+\frac{\sqrt{2\sigma}}{A_0X}\xi\left(\tau\right) \\ \frac{d\tilde{R}}{d\tau} &= \frac{l_R}{hR_0R_0}-\frac{m_R}{hR_0}\tilde{R}+\frac{\beta\frac{l_A}{hR_0}}{hR_0}\tilde{A}\tilde{R}\end{aligned}$$

$$\frac{d\tilde{A}}{d\tau} = 1 - \frac{m_A}{hR_0}\tilde{A} + \frac{\gamma A_0}{hR_0}\tilde{A}^2\left(1 - \frac{A_0\tilde{A}}{C}\right) - \tilde{A}\tilde{R} + \frac{\sqrt{2\sigma}}{A_0 hR_0}\xi(\tau)$$
$$\frac{d\tilde{R}}{d\tau} = \frac{m_R}{hR_0} - \frac{m_R}{hR_0}\tilde{R} + \frac{\beta A_0}{hR_0}\tilde{A}\tilde{R}$$

we can now define the following dimensionless parameters:

$$\begin{aligned} G &= \frac{\gamma A_0}{hR_0} \\ D &= \frac{m_R}{hR_0} < 1 \\ C &= \frac{C}{A_0} \\ B &= \frac{\beta A_0}{hR_0} = \frac{\beta l_A}{hR_0^2} \\ \Sigma &= \frac{\sigma}{A_0^2 h^2 R_0^2} \end{aligned}$$

and result with the folllowing dimensionless equations (denoting $\tilde{R} = R$, $\tilde{A} = A, \tau = t$)

$$\begin{aligned} \frac{dA}{dt} &= 1 - AR + GA^2\left(1 - \frac{A}{C}\right) \\ \frac{dR}{dt} &= D(1 - R) + BAR \end{aligned}$$

In the case we're considering $C \gg 1$, since each flare exceeds the steady state by a large amount. Furthermore we assume that $B$ is small, to match the observation that regulatory T cells have a powerful inhibitory effect despite having a relatively low abundance relative to the effector cells.

##### 1.3 Nullclines and Steady State

Nullclines:

$$\begin{aligned} R &= \frac{1 + GA^2\left(1 - \frac{A}{C}\right)}{A}, \qquad \text{A nullcline} \\ R &= \frac{D}{D - BA}, \qquad \text{R nullcline} \end{aligned}$$

The extrema of the A nullcline (assuming that there are - what is the condition for that?). Deriving the A nullcline:

$$\frac{\partial R_{\text{A nullcline}}}{\partial A} = \frac{-2A^3G + A^2CG - C}{A^2C} = 0$$

to solve we consider approximate two regimes. The first is far from the acrrying capacity $A \ll C$:

$$\begin{aligned} 0 &= A^2G - 1 \\ A_{min} &= \frac{1}{\sqrt{G}} \end{aligned}$$

$$
\begin{aligned}
R_{min} &= \left.\frac{1+GA^2\left(1-\frac{A}{C}\right)}{A}\right|_{A_{min}} \\
&\approx 2\sqrt{G}
\end{aligned}
$$

other regime is close to the carrying capacity $A \to C \gg 1$:

$$
\begin{aligned}
A_{max} &= \frac{C}{2} \\
R_{max} &= \frac{2\left(\frac{GC^2}{8}+1\right)}{C} \\
&\approx \frac{GC}{4}
\end{aligned}
$$

Jacobian:

$$
\begin{aligned}
J &= \begin{pmatrix} \frac{\partial \dot{R}}{\partial R} & \frac{\partial \dot{R}}{\partial A} \\ \frac{\partial \dot{A}}{\partial R} & \frac{\partial \dot{A}}{\partial A} \end{pmatrix} \\
&= \begin{pmatrix} \frac{\partial}{\partial R}\left[D-DR+BAR\right] & \frac{\partial}{\partial A}\left[D-DR+BAR\right] \\ \frac{\partial}{\partial R}\left[1-AR+GA^2\left(1-\frac{A}{C}\right)\right] & \frac{\partial}{\partial A}\left[1-AR+GA^2\left(1-\frac{A}{C}\right)\right] \end{pmatrix} \\
&= \begin{pmatrix} -D+BA & BR \\ -A & -\frac{A^2G}{C}+2AG\left(1-\frac{A}{C}\right)-R \end{pmatrix}
\end{aligned}
$$

Steady state: we assume that $B$ is small so the steady state of R is $R_{st}=1$. We find

$$
\begin{aligned}
A_{st} &= \frac{\frac{l_A}{hR_0}}{A_0} = 1 \\
A_{th} &= \frac{\frac{hR_0+hR_0\sqrt{1-\frac{4\gamma l_A}{hR_0}}}{2\gamma}}{A_0} \\
&\approx \frac{hR_0}{\gamma A_0} = \frac{1}{G}
\end{aligned}
$$

#### 1.4 Flare Ups - Dimensional

We begin by re-writing the equations in their dimensional form:

$$
\begin{aligned}
\frac{dA}{dt} &= l_A - m_A A + \gamma A^2\left(1-\frac{A}{C}\right) - hAR \\
\frac{dR}{dt} &= l_R - m_R R + \beta AR
\end{aligned}
$$

First of all we want to find the steady state. The nullclines are:

$$
\begin{aligned}
R &= \frac{l_A - m_A A + \gamma A^2\left(1-\frac{A}{C}\right)}{hA} \\
R &= \frac{l_R}{m_R - \beta A}
\end{aligned}
$$

we approximate the leftmost part of the A nullcline, on which the steady state resides, as:

$$R \approx \frac{l_A}{hA}$$

solving for the steady state:

$$\begin{aligned} R_0 &= \frac{l_A}{hA} \\ A_{st} &= \frac{l_A}{hR_0}\left[\frac{c^1 t^{-1}}{c^{-1}t^{-1}c} = c\right] \end{aligned}$$

and the steady state of R is approximately $R_{st} \approx R_0$.

Next we define the effective potential of the A equation:

$$\begin{aligned} \frac{dA}{dt} &= l_A - m_A A + \gamma A^2\left(1 - \frac{A}{C}\right) - hAR + \sqrt{2\sigma}\xi(t) \\ &\approx l_A - (m_A + hR_0)A + \gamma A^2 + \sqrt{2\sigma}\xi(t) \\ &= -\frac{\partial U}{\partial A} + \sqrt{2\sigma}\xi(t) \end{aligned}$$

with

$$U(A) = -l_A A + \frac{m_A + hR_0}{2}A^2 - \frac{\gamma}{3}A^3$$

To find the rate of flares , we need to compute the minima and maxima of this potential. These are given by the zeros of $\frac{\partial U}{\partial A}$.

$$\begin{aligned} l_A - (m_A + hR_0)A + \gamma A^2 &= 0 \\ A_{st} &= \frac{hR_0 + m_A - \sqrt{(hR_0 + m_A)^2 - 4\gamma l_A}}{2\gamma} \\ A_{th} &= \frac{hR_0 + m_A + \sqrt{(hR_0 + m_A)^2 - 4\gamma l_A}}{2\gamma} \end{aligned}$$

where $A_{st}$ is the local minima of the potential and $A_{th}$ is the local maxima. To use Arrhenius/Kramers approx. we need to compute:

$$\begin{aligned} \Delta U &= U(A_{th}) - U(A_{st}) \\ &= \frac{\left((hR_0 + m_A)^2 - 4\gamma l_A\right)^{3/2}}{6\gamma^2} \\ &= \frac{(hR_0 + m_A)^3\left(1 - \frac{4\gamma l_A}{(hR_0 + m_A)^2}\right)^{3/2}}{6\gamma^2} \\ &= \frac{(hR_0 + m_A)^3\left(1 - \frac{6\gamma l_A}{(hR_0 + m_A)^2}\right)}{6\gamma^2}, \gamma l_A \ll (hR_0 + m_A)^2 \\ &= \frac{(hR_0 + m_A)^3}{6\gamma^2} - \frac{(hR_0 + m_A)\,l_A}{\gamma}, m_A \ll hR_0 \end{aligned}$$

$$
\begin{aligned}
&\approx \frac{(hR_0)^3}{6\gamma^2} - \frac{(hR_0)\, l_A}{\gamma}, \text{proxying } A_0 = \frac{l_A}{hR_0} \\
&= \frac{(hR_0)^3}{6\gamma^2} - \frac{(hR_0)^2 A_0}{\gamma}
\end{aligned}
$$

As well as calculating the second derivative of the potential in these locations:

$$
\begin{aligned}
U''(A) &= hR_0 - 2\gamma A \\
U(A_{th}) &= -hR_0 \\
U''(A_{st}) &= hR_0 - \frac{2\gamma l_A}{hR_0} \approx hR_0
\end{aligned}
$$

So the Kramer approximation yields:

$$
\begin{aligned}
\langle T \rangle &= \sqrt{U''(A_{th})\, U''(A_{st})} \exp\left[\frac{\Delta U}{\sigma}\right] \\
&= hR_0 \exp\left[\frac{(hR_0)^3}{6\gamma^2 \sigma} - \frac{(hR_0)\, l_A}{\gamma}\right]
\end{aligned}
$$

Therefore the average relapse rate is:

$$
\begin{aligned}
\langle r \rangle &= \frac{1}{hR_0} \exp\left[-\frac{(hR_0)^3}{6\gamma^2 \sigma} + \frac{(hR_0)\, l_A}{\gamma}\right] \\
&\approx \exp\left[-\frac{(hR_0)^3}{6\gamma^2 \sigma} + \frac{(hR_0)\, l_A}{\gamma}\right]
\end{aligned}
$$

#### 2 Effect of Parameters on the Shape of Flares

In the following figure, we show graphically the effect of each parameter on the shape of the flares.

Figure 1: **Effect of different parameters on the shape of flares.**The parameters used were: $B = 0.001, G = 0.25, D = 0.15, C = 1000$

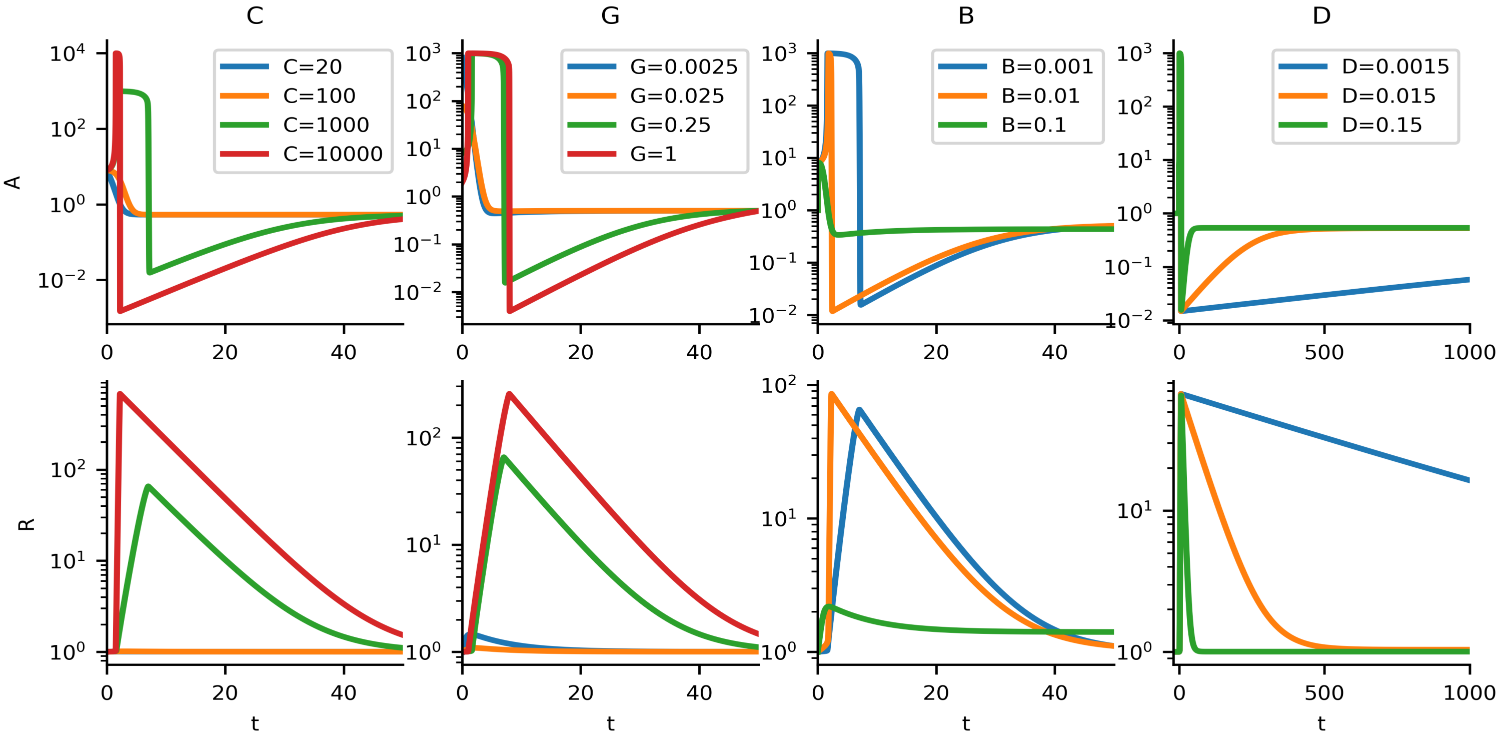


It can be seen that increasing the carrying capacity C allows for higher peaks in both A and R. Increasing G results both in higher peaks of A and R, and also longer time of active inflammation (high A) and longer refractory period. Increasing B, however, results in lower peaks of A and R and shorter active inflammation periods. The main effect of increasing D is decreasing the refractory period (high R, low A).

### 3 Justification for noise on the A equation only

As we mentioned, in order to have effective shutdown of the flare, we must have $\beta \ll h$ or equivalently $B \ll 1$.This results in nearly horizontal vector field near the steady state (as can be seen in Fig 2A,B in the main text). This means that fluctuations in the vertical direction (i.e, in the R equation) will have a lesser impact on the chance to cross the threshold.

We note here that since the differential equations don't exhibit a potential $U(A, R)$ that will satisfy both:

$$\begin{aligned} \frac{\partial U}{\partial A} &= -\frac{dA}{dt} \\ \frac{\partial U}{\partial R} &= -\frac{dR}{dt} \end{aligned}$$

and hence introducing noise to both equations prohibits analytical analysis using a Kramers approach.

Taking these two considerations into account, we decided to employ noise only on the A axis, to be able to obtain meaningful and comprehensive analytical results.

### 4 Additional Phase portraits

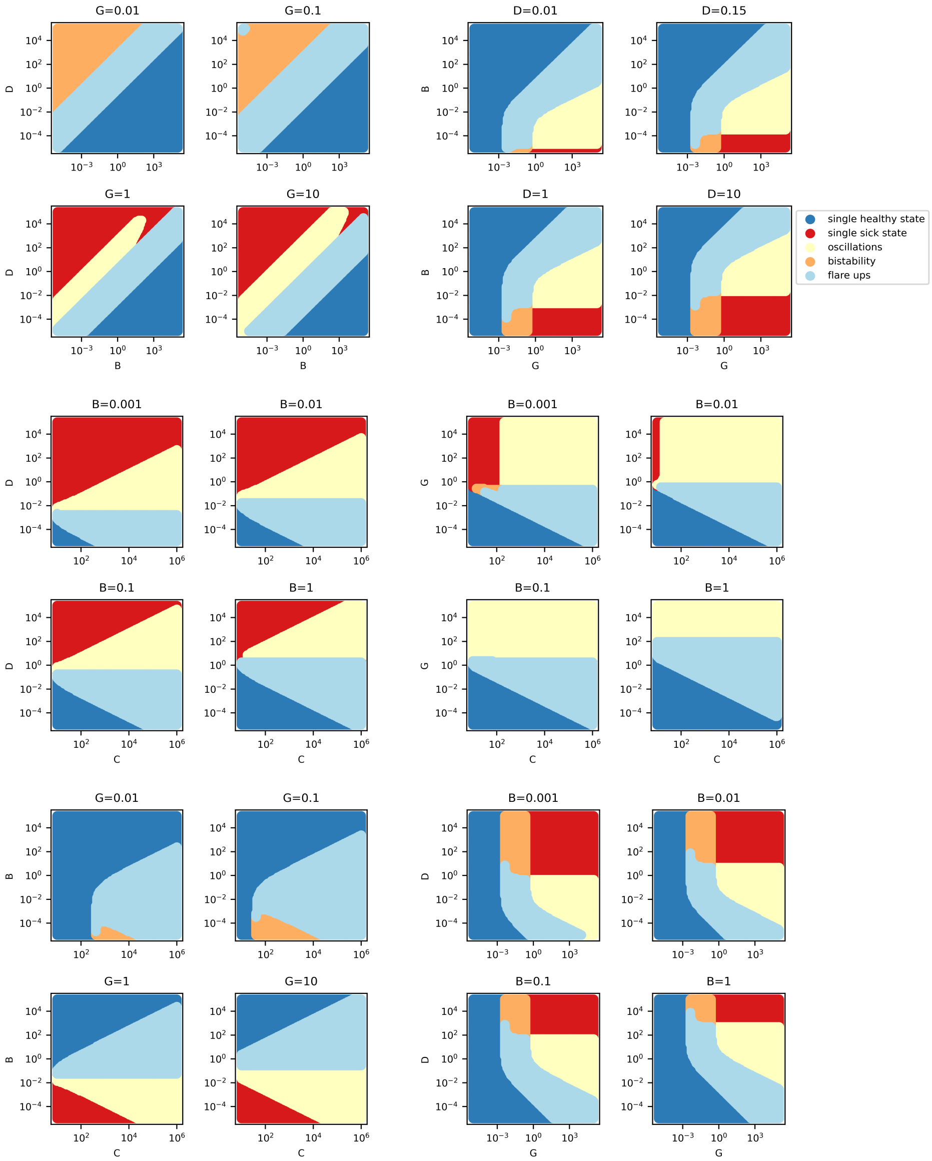

### 5 Other Models

We studied alternative models as described next.

#### 5.1 Inhibition of A by dividing by R

This model bases of the same variables as the model presented in the main text, but encompasses a different form of interaction between the auto-reactive T cells (A) and the regulatory T-cells (R), in which R inhibits the proliferation rate by dividing the auto-stimulation prolfieration term, and A raises the prolifetion rate of R in a linear fashion The model equations are:

$$\begin{aligned}\frac{dA}{dt} &= l_A - m_A A + \gamma\frac{A^2}{R}\left(1 - \frac{A}{C}\right)\\ \frac{dR}{dt} &= l_R - m_R R + \beta A\end{aligned}$$

where the parameters have the same biological meaning (even though different units) as those presented in the main text.

Although the nullclines in this model do posses the classical N-shaped form that allows for excitability, the range of parameters needed in this form did not result in strong spikes with high amplitude of A.

#### 5.2 Dynamics limited by healhty tissue

This model describe dynamics of different variables than the one presented in the main text: auto-reactive cells (A) and healthy tissue (H). The model equations are:

$$\begin{aligned}\frac{dH}{dt} &= l_H - m_H H - \beta H A\\ \frac{dA}{dt} &= l_A - m_A A + \beta\gamma H A^2\left(1 - \frac{A}{C}\right)\end{aligned}$$

With:

- $l_H\left[\frac{cells}{time}\right]$ : production rate of the healthy tissue cells
- $m_H\left[\frac{1}{time}\right]$ : natural mortality rate of the healthy tissue cells. $\frac{1}{m_H}$ is the turnover time of the healthy tissue. $\frac{l_H}{m_H}$ is the steady state of the healthy tissue without auto-immune killing ($\beta = 0$).
- $\beta\left[\frac{1}{time\cdot cells}\right]$ : the killing rate.
- $l_A, m_A$ are the same as $l_H, m_H$ but for the auto-immune cells.
- $\gamma\left[\frac{1}{cells}\right]$ : the response of the immune system to the killing
- $C\,[cells]$ : the carrying capacity of the immune system.

In this model an intrinsic separation of timescale is present, since the typical turnover of the auto-reactive cells is of the order of hours to days, whereas the typical turnover rate of the tissue is of the order of weeks to months. This separation of timescales allows for analysis similar to the one done in the FitzHugh-Nagumo model.

However, in this model, the flare up is shut off when the healthy tissue level declines close to zero. This stands in contrast to the biological observation that regulatory T cells are mainly responsible for the shutdown of the flare. Moreover, in MS and other diseases, the healthy tissue (e.g, in MS, the total level of myelination) does not decline in such a drastic manner during a flare up of a disease, and therefore this model was rejected.
